# Molecular Evolution and Functional Divergence of Heterodisulfide Reductase A (HdrA)

**DOI:** 10.1101/2024.09.26.615249

**Authors:** Mousumi Banerjee

## Abstract

Heterodisulfide reductase is a key enzyme in methanogenesis pathway in archaea, which helps to recycle the coenzymes after methane is formed from C1 carbon sources. Homologous enzymes are observed in non-methanogenic archaea and prokaryotes but their functional and evolutionary details are not fully explored. This report aims at understanding the distribution of one such homologous enzyme, heterodisulfide reductase A (HdrA); it’s domain architecture, overall structural contribution in biological function and evolutionary details in organisms other than methanogen. Exhaustive sequence analysis from both archaea and prokaryotic lineages reveals the presence of highly conserved homologues which probably have an altered oxido-reductase function. Phylogenetic analysis and co-clustering showed the differential diversity of the homologues enzyme in metabolically diverse groups. The distribution of Heterodisulfide homologues in both archaea and prokaryotes shows differential expansion of this protein by multiple horizontal gene transfer events. Conserved domain analysis and structural modelling of different phylogenetic clades shows sequence and structural diversity at the C-terminal domain. Bimolecular network analysis highlights the functional association and interactions of the heterodisulfide homologues with diverse protein partners which participate in different metabolic function. HdrA involved in species specific metabolic function while the functional conservation is evident between close related species. Analysis of structural domain within HdrA’s reveals that clade specific sequence which helps to differentiate structural domain arrangement and contributes to functional constrains. The presence of HdrA protein in methanogenic archaea and non-methanogens suggests that the protein function involving redox reaction and electron transfer shared a common history.

## Introduction

Methanogens are the methane producing archaea contributes 1Gt of methane every year with a net high estimates ~ 75% of the global methane emission annually (**1,2**). Methanogenic archaea are important for their metabolic product, methane, which makes 25 times more contribution than the greenhouses gas carbon dioxide to the earths atmosphere (**3,4**). These anaerobic methane producing archaea maintain an exotic metabolic pathway to produce methane from C1 carbon sources with the help of extraordinary enzymes and co-factors. Studies on methanogenesis or methane metabolism showed how this microbial world conserve energy to grow and helped to understand the bioenergetics and thermodynamic basis of life (**5–7**). However, complete understanding of biodiversity of these exotic enzymes and evolution are rare. Heterodisulfide reductase, the last enzyme in the methanogenesis pathway of archaea, is a multi-domain enzyme complex, exhibits multifunctional activity, including hydrogenation, thiol-disulphide reduction, electron bifurcation and electron transfer (**8–10**). The critical oxido-reductase enzyme in methanogenesis pathway known for its energy conservation yet their functional and evolutionary details are not fully explored. Heterodisulfide reductase enzyme complex is the last enzyme in methanogenesis pathwaywhich catalyse the thiol disulphide conversion of disulphide bonded Coenzyme B (CoB) andCoenzyme M (CoM) [CoMS-SCoB] to its individual components and plays crucial role to recycling the rare CoMSH and CoBSH into the pathway while completing the methane production cycle fromone carbon sources. (**11–14**). The subunits poses Flavin moiety Fe-S cluster core and to perform the catalysts reaction and conserve energy through flavin based electron bifurcation (FBEB) where [Fe-S] helps in electron transfer. (**15,16**). Heterodisulfide reductase plays a key role in the energy metabolism of methanogenic archaea where it functions as the terminal electron acceptor of an energy-conserving electron transport chain (**12,16**). The reducing equivalents required for the reduction are provided either by H_2_ or by reduced coenzymes. Heterodisulfide reductase (Hdr) generally works as a multi subunit complex like HdrABC, when present in the cytosol or HdrDE, while embedded in the cell membrane. The HdrABC complex catalyses an Fe-S cluster assisted disulfide reduction reaction which is integrated into a flavin-based electron bifurcation (FBEB) process, a mode of energy coupling which optimises the energy yield of the cell (**15, 17–22**). The key subunits are HdrA, which carries the electron-bifurcating flavin adenine di-nucleotide (FAD), and HdrB, which has been proposed to be the heterodisulfide reductase site (**15**). HdrA, the FAD binding domain which also carries electron bifurcation reaction, homologs are found in many other microorganisms i.e., anaerobic methanotrophic archaea (**23**), sulfate-reducing bacteria (**24**) and archaea (**25**), sulfur-oxidizing bacteria (**26**), acetogenic bacteria (**27**), and metal-reducing bacteria (**28**). Although most remain biochemically uncharacterised, the electron bi-furcation modules are assumed to be connected. Specific roles of HDrA protein in different organisms are reportedhowever evolution of a specific enzyme which were supposed to belong only in methanogenic pathway is not studied.

Understanding functional diversities in enzyme is an expanding area in evolutionary biology. How ancestral sequences and scaffolds diversified to give rise different enzyme functionsis one of the important question in enzyme evolution to ask. Here, we report a study of an archaea specific metabolic enzyme Heterodisulfide reductase (Hdr) and its sequence and structural homologues to understand how the enzyme evolved and diversified over 2 billion years. A completehomologous sequence analysis followed by phylogeny reveals how the HdrA sequence is distributed and diverge in methanogenic archaea, non-methanogenic archaea and bacterial species. Conserved domain search and structural analysis reflects how the fold and domain preservation related to the sequence divergence and finally the protein protein network analysis of the homologous sequences explains how the sequence diversity translate to metabolic diversity preserving the ancient HdrA protein and repurposing them to achieve metabolic diversity and sustainability.

The early earth anaerobes are very difficultly to culture under laboratory environment thus an in-silico approach were followed to analyse large set of sequences and their structure to find out evolutionary relationship and the possible mechanism of energy conservation. The present analysis of HdrA sequences aimed at understanding the distribution, phylogeny, domain architecture, conservation, expansion and evolution of biological function in geo-biologically relevant sequences. Extensive analysis of homologous HDR sequences from archaeal and bacterial lineages reveals that the sequences are highly conserved, cysteine rich, contain low complexity repeats and unnatural amino acids. It was curios that if HdrA is present in various different organisms other than archaea whether its function is preserved, how methanogens, so early in the evolution, solved the complex problem of redox regulation and energy conservation through electron bifurcation. We also wanted to study the evolution of HdrA in early earth organisms and how the sequences were diverged. Phylogenetic clustering of related sequences from methanogens and non-methanogenic archaea and bacteria reveals that the evolution of similar sequences, through horizontal gene transfer (HGT), with a similar redox function for the respective redox equivalents. Domain conservation and structural analysis reflects the preservation of heterodisulfide reductase equivalent entity for electron transfer and energy conservation through electron bifurcation is the key to survive and thrive in early anaerobes.

Our understanding of different HDR sequences from conserved domain analysis and structural modelling advanced our insight into the conformation specific regulation of their distinct function. The structure of the HDR C-terminal domain their structural and functional interplay with the sequences surrounding these domains and discuss a potential evolutionary path.

## Materials and Methods

### Sequence search and similarity analysis

The initial sequence search with *M. jannaschii* Heterodisulfide A sequence (Uniprot id : P60200) was performed in Uniprot. [https://www.uniprot.org] later the sequences were obtained from National Centre for Biotechnology Information (NCBI, https://blast.ncbi.nlm.nih.gov) withnon redundant (nr) protein dataset using the Position-Specific Iterative Basic Local Alignment Search Tool (PSI-BLAST) search (**30**) with *M. jannaschii* HDR sequence (Uniprot id P60200) asthe query sequence with an E-value cutoff of 0.0001 and default parameter settings. PSI-BLAST iterations were performed until no new BLAST hit was retrieved. A final set of 500 sequences from non-redundant sequences were chosen for further analysis. In order to detect the domain boundaries within the sequence, Conserved Domain search was performed against the NCBI Conserved Domain Database (**31**) for the query sequence.

### Multiple sequence analysis

Protein sequences were aligned using Clustal Omega (**32**). The poorly aligned sequences were then manually inspected and adjusted for the gap penalty.

### Phylogeny

Phylogenetic analyses were performed with the MEGA package Version 11.0 (**33**). First, protein sequences of Hdr were aligned with the MUSCLE program implemented in the MEGA package with up to 100 iterations. Obtained alignment was subjected to maximum Likelihood (ML) method of phylogeny reconstruction available in the MEGA 11.0 package. For ML 1000 bootstraps were applied. Obtained phylogenetic trees were depicted with the Tree Explorer program of the MEGA package. The percentage of replicate trees in which the associated sequences cluster together in the bootstrap test (1000 replicates) were calculated, and branches with 50% bootstrap cutoff were collapsed. The evolutionary distances were computed using the Poisson correction method and are in the units of number of amino acid substitutions per site. All positions containing alignment gaps and missing data were eliminated only in pair-wise sequence comparisons (pairwise deletion option).

### Conserved domain analysis

All the gene products, containing at least one Hdr domain, were then searched against the protein conserved database CDD (**31**) at an E-value of 10^−2^, to identify and classify coexisting domains. Protein domain identification is limited by the sensitivity of the search and the diversity of protein domains in the database. CDD was used to increase both the sensitivity and the protein diversity.

### Structure analysis and protein structure modelling

The conserved residues were mapped onto the HdrA structure from *M. thermolithotropicum* (PDB ID:) also the domain conservation was mapped onto the same PDB structure to visualise the difference clearly using PyMOL (**34**). 11 different homologous sequences were modelled using SWISS-MODEL (https://swissmodel.expasy.org) (**35**). Structures were validated using PROCHECK software (**36**). The model with a highest confidence was chosen for further study.

### STRING analysis

A comprehensive protein-protein interaction network analysis was done using STRING database (version 11.5) (**37**) to predict PPIs which can be applied to predict functional interaction of proteins. Species specific searches were done for individual sequences, where an interaction score were set >0.7 to construct individual PPI network.

## Results

### Sequence query

Heterodisulfide reductase is a multi domain protein which exists as a HdrABC or HdrDE complex (**38,39**). Here HdrA sequence from *M. jannaschii* is taken as a standard (Uniprot id : P60200) to analysed homologous HdrA sequences from archaea and other kingdoms. The HdrA sequence is ~600-700 amino acid long and contains a pyridine nucleotide disulphide oxido-reductase domain and a ferridoxin domain. The sequences have 2 low complexity regions (between 142-162 aa and 615-630aa; the amino acid numbering is according to *M. jannaschii*) and 2 internal repeat regions (256-323 aa and 543-614 aa) (**Figure 1**). The natural sequences are cysteine and glycine rich however, a few archaea sequences including *M. jannaschii* contains unnatural amino acid selenocysteine. Although Hdr is known for its role in methanogenesis in archaea, homologous sequence search using PSI-BLAST shows a wide distribution of this particular protein sequence from archaea to bacteria including euryarcheotes, CFB group of bacteria, GNS bacteria and d-proteobacteria (**Figure 2**). However, no eukaryotic sequences was found using the present specific search criteria (PSI-BLAST Materials and methods). **Figure 2** sows the distribution of 500 HDr sequence data in different organisms. There are total seven subgroups where the significant occupancies are; archaea 32%, euryarchaeots 21%, bacteria 18% and D-proteobacteria 15%. No eukaryotes were found in the final dataset where 40% and above sequence identity cut off were set.

**Figure 1:**
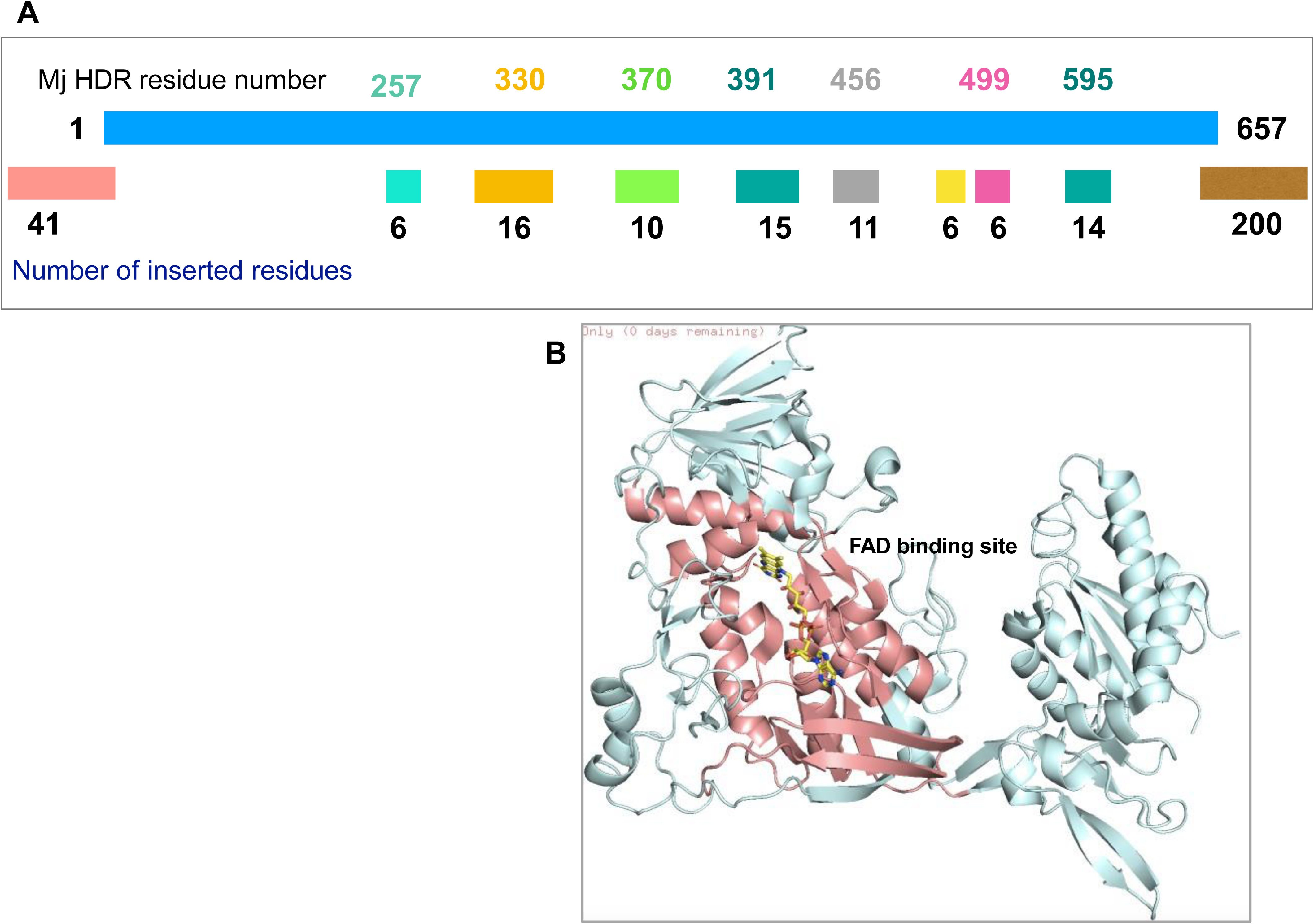
The amino acid sequence and the 3D structure of HdrA protein. (A) The highlighted residues in xx are FAD binding domain, variable domain, Ferredoxin binding domain and C-terminal domain. Fe-S binding motifs are shown in bold letters. (B) Different domains of HdrA in 3D and the FAD binding site.

**Figure 2:**
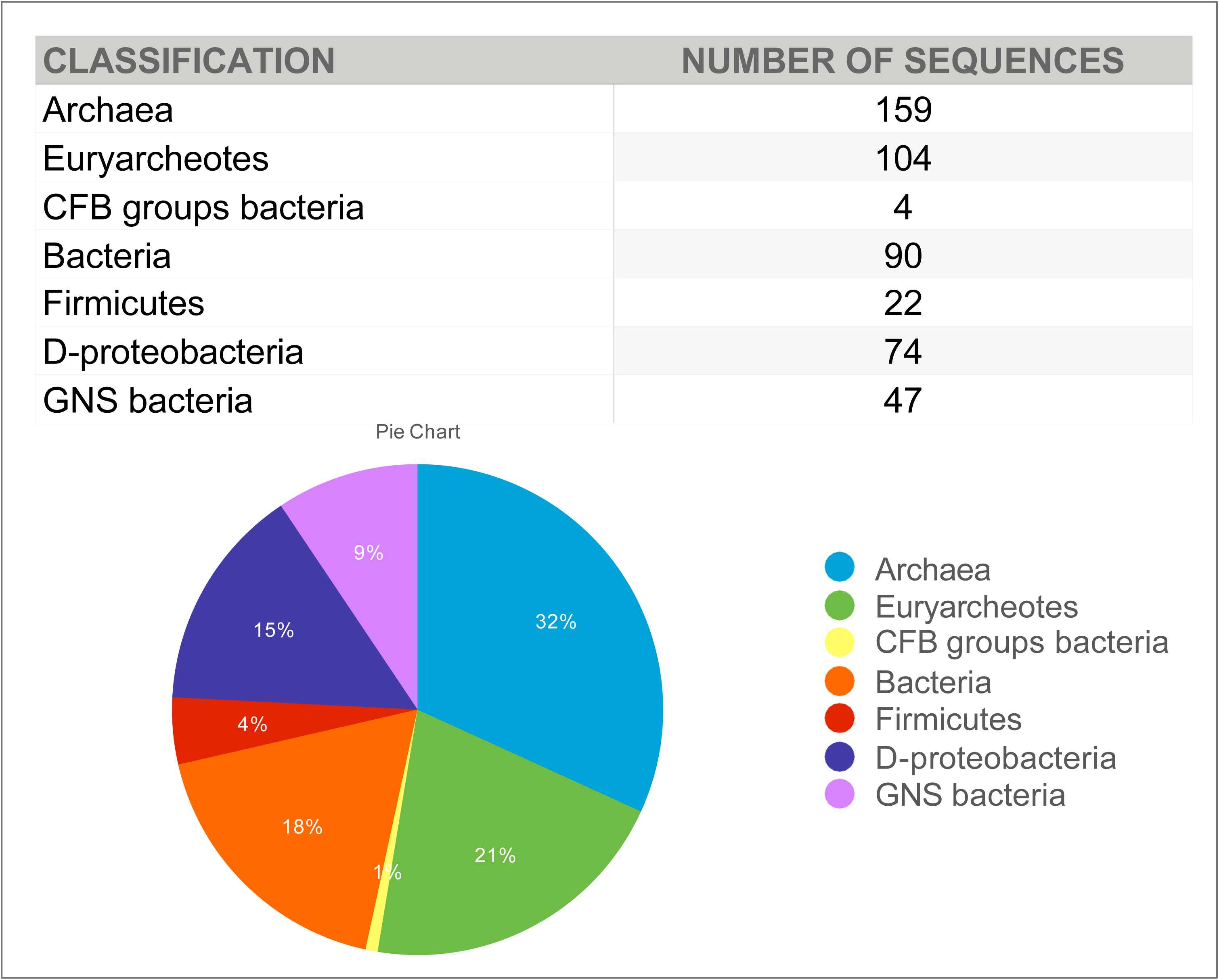
Species specific classification and distribution of homologous HdrA sequences in prokaryotes and archaea lineages.

### Multiple sequence alignment

Multiple sequence alignment of 500 HDrA sequences were done using CLUSTAL omega (**32**). Quality of the alignment was verified by the overall score of the alignment. The sequence length of the HDrA sequence dataset varied from 640-775 and overall pairwise sequence identity is given in Table S1 in supplementary section. There are 39 residues out of an average 700 residue protein are conserved, which is ~5.5% which is very high. List of conserved position, Sequence repeats, Amino acid repeats, sequence insertion, sequence deletion are shown in Figure 1. The Figure 1 shows that there are five major classes of organisms from where the major differences in the alignment are coming they are Syntrophomonus (anaerobic non phototropic bcterium), Desulfobacteria (mesophilic, autotrophic, sulphate reducing bacteria), Actinobacteria (shares features with both bacteria and fungi and play role on recycling biomaterial), Syntrophic archeaum and Armatimonadota (aerobic, gram negative bacterium). HDrA has around 17 conserved cystine residues with six highly conserved CXC, CXXCXXCX(3)C, CXXC, CXXXCC, CX(6)CXXC, CXXC motifs for Fe-S binding. The HDrA sequence has two extremely variance regions between residues 450-500 and 590-650.

### Maximum likelihood phylogeny

The evolutionary history was inferred by using the Maximum Likelihood method and JTT matrix-based model (**33**). The tree with the highest log likelihood (−223433.92) is shown. Initial tree(s) for the heuristic search were obtained automatically by applying Neighbor-Join and BioNJ algorithms to a matrix of pairwise distances estimated using the JTT model, and then selecting the topology with superior log likelihood value. This analysis involved 501 amino acid sequences. There were a total of 1072 positions in the final dataset. Evolutionary analyses were conducted in MEGA X (33) (Figure 3).

**Figure 3:**
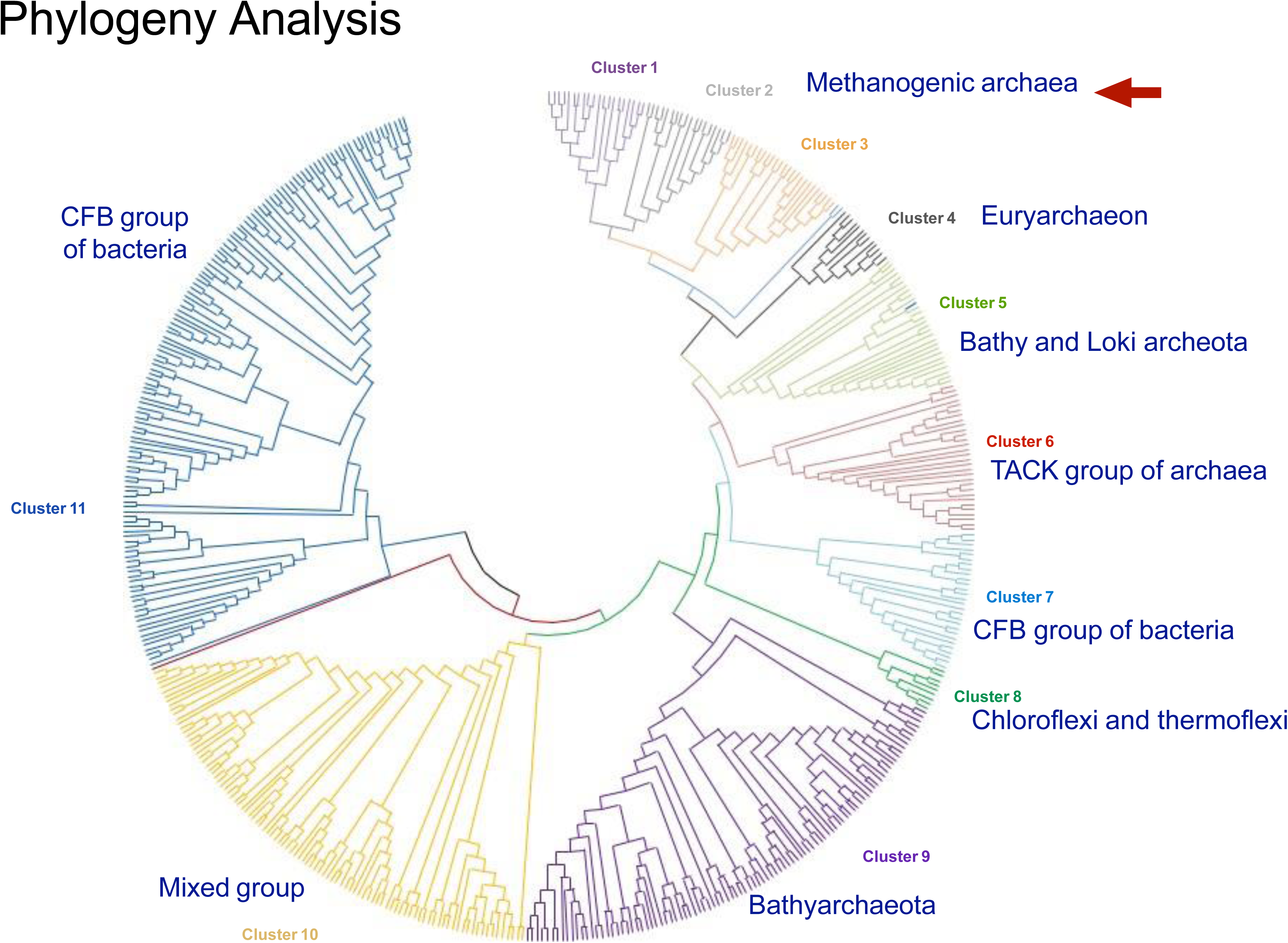
Maximum likelihood phylogenetic tree of HdrA homologues from archaea and bacteria. The super phyla are highlighted with colour ranges. The tree is constructed using MEGA 11.0 packages and 1000 bootstrap were calculated and branches of >50% value are taken. The average bootstraps support values for each phylum is provided in the supplementary section.

The HdrA sequences differ significantly from one another in their sequence length (Figure 1). Pairwise sequence comparison studies indicate that the overall sequence similarity between members of different sequences is in the range of 40-48% and much of the observed similarity is seen around the conserved residue stretches. A phylogenetic tree was constructed based on Maximum Likelihood method (see Methods) to investigate the evolutionary relationship among homologous sequences (**Figure 3**). All Hdr related sequences were diversified/ classified in 11 different major clades (**Figure 3**). Each clade contains between 15-30 members. Clade 1-3 in**Figure 3** showed mostly methanogenic archaea containing HDrA enzyme with various subgroups; Clade 4 is a mixed clade the organisms shows anaerobic methane oxidation, Clade 5 is mostly contains bathyarchaeota and lokiarcheota mixed clade, Clade 6 contains mostly very and korarcheota group, clade 7 is made up of all anaerobic bacteria including formictes, actinobacteria, cholroflexi etc, clade 8 contains thermoflexi bacteria but presence of chloroflexi is also evident, clade 9 exclusively shows Bathyarchaeota, Clade 10 is again a mixed clade −11 are mostly bacterialbut this are aerobic class. As shown in **Figure 3** phylogeny clade 1-3 encompasses all known methanogens and represents themselves in monophyletic lineage at the exclusion of other archaea groups. Clade 10 and 11 showed early diversion as well as a mixed clade identity. However, the pairwise sequence comparison of Clade 10 and 11 with the Methanogen sequences showed 70-75% sequence identity which is significant compared to Clade 6/7 where the sequence identity is 40-45%. Yet these lineages (Clade 10 and 11) are distantly related as evidenced by their survival in aerobic condition. Strict “methanogens” including methanobacterials, methanolpyrales, methanococcales, methanomicrobials and methanosarcinales shows monophyly (**Figure 3**) which indicates their evolutionary ancestry as well as their similarity of methane production with respectto HDrA enzyme. However, among prokaryotes and archaea lineages like bathyarcheota, euryarcheota etc showed patchy appearances in the phylogenetic tree.

### Conserved domain (CDART) analysis

The conserved domain architecture retrieval were performed on the curated 501 sequence data set using similarity searches (**31**). CDART finds protein similarities across significant evolutionary distances using sensitive protein domain profiles rather than by direct sequence similarity. Searches can be further refined by taxonomy and by selecting domains of interest. HdrA is a polyferrodoxin belongs to heterodisulfide reductase superfamily which contains beta-alpha-beta-beta-alpha-beta fold. HdrA, which is prevalent in methanogenic archaea however, other archaea and bacterial homologous are also investigated. The core sequence belong to HDR superfamily but the N and C terminal ends often have diversesequences and folds. **Figure 4** shows the sequence alignment of 11 different clades and their domain arrangement. The variation of domain arrangement and appearance of new domains along with core polyferrodoxin domain is observed. The domains are mostly different at theN or C terminal end of the core protein. Clade specific domain searches for the corresponding HdrA sequences reveals that the strict methanogens which belongs to clade 1 and 2 mostlyhost the core polyferrodoxin domain where other clades introduces NAD binding, FlpD, Fer4, Fer7, Fer9 and Pyr_Rdx domains in various different combinations (**Figure 5**).

**Figure 4:**
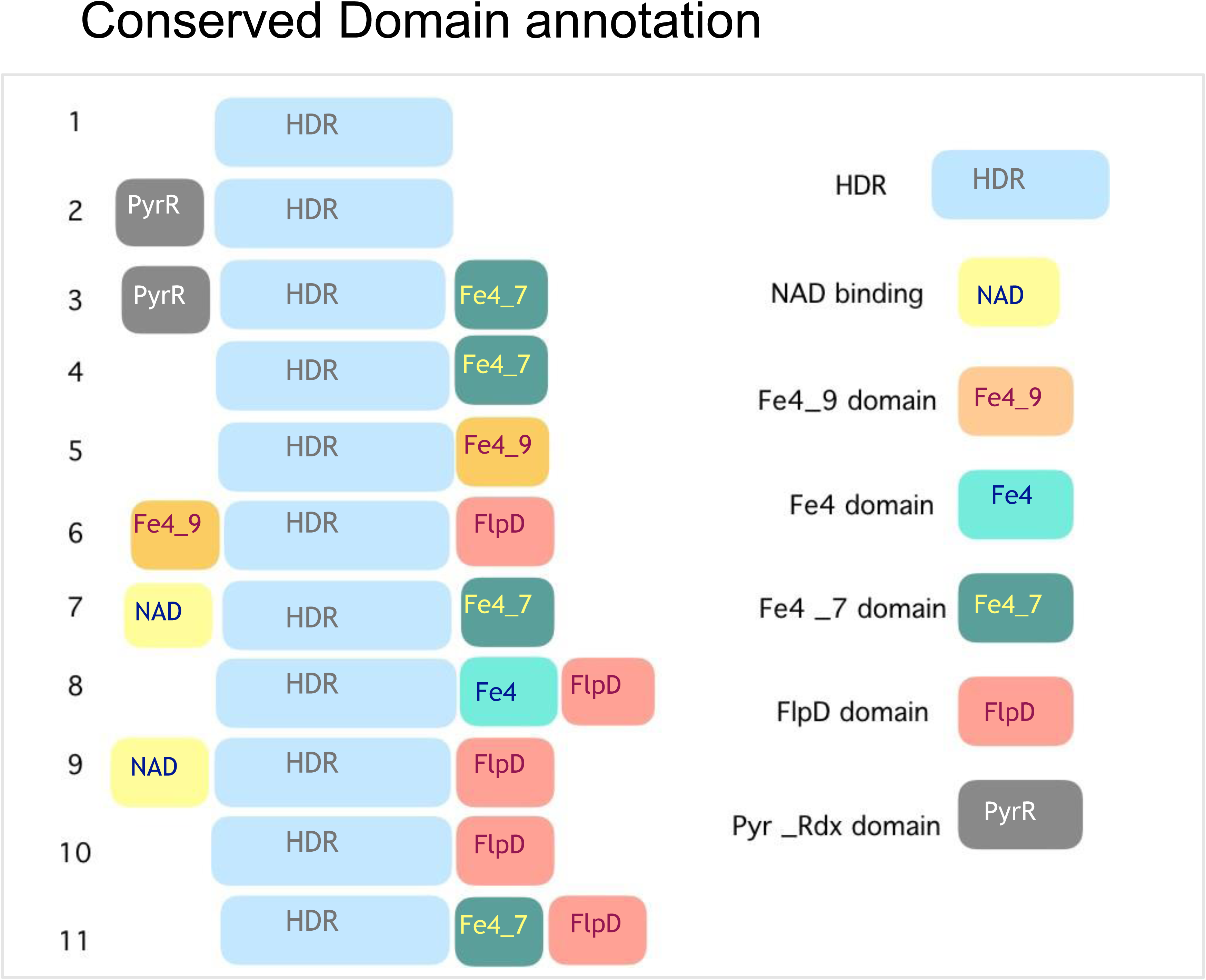
Graphical representation of domain architecture present in HdrA homologues. HDR-heterodisulfidereductase core domain in (pale blue), NAD-NAD binding domain (Yellow), Fe4_9-ferredoxin domain (orange), Fe4-Fe4S4 alpha helical domain (teal green), Fe4_7-Ferredoxin binding dominion (green), FlpD-Methylviologen reducing hydrogenase domain (pale red), Pyr-Pyr_rdx domain (grey).

**Figure 5:**
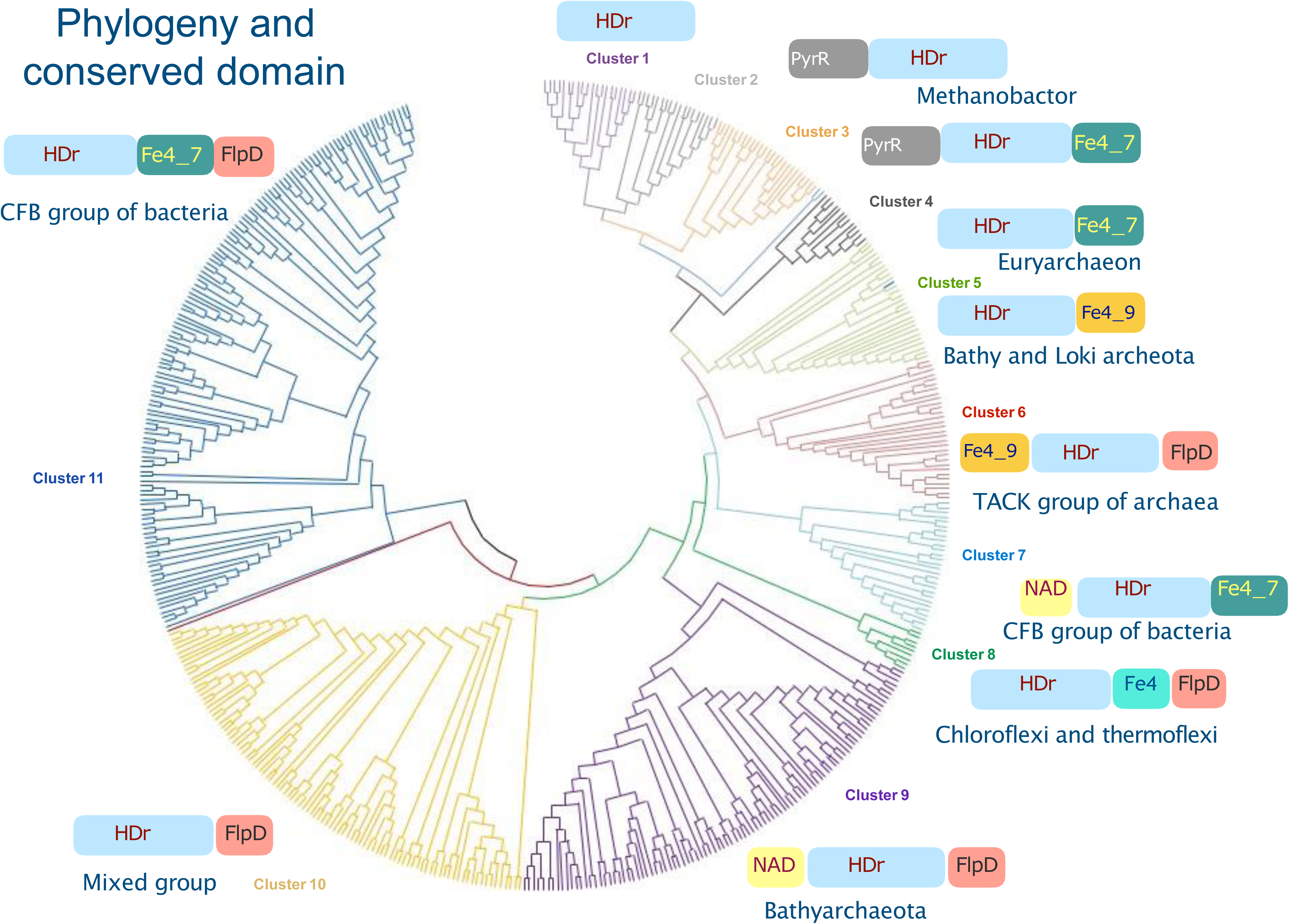
Domain architecture of HdrA homologues mapped onto sequence phylogeny tree.

### Structure and C-terminal domain

HdrA structure contains N-terminal domain (residues 1-135), a flavin binding domain (residues 145-236; 315-567), an inserted ferredoxin domain (residues 270-320) and a C-terminal ferredoxin domain (residues 570-625) (**Figure 6**). (PDB id: 5ODH) (**15**). Figure xx presents the modelled HdrA structures from 11 different representative homologous sequences of different phylogenetic clade. The overall structures remained similar with 4 different domain organisation however, the inserted ferredoxin and the C-terminal ferredoxin domain showed differences. The ferredoxin inserted domain is mostly made up of loops and coiled structure where as the C-terminal ferredoxin domain have two shorter helices and two pairs of beta sheet connected by loop structures. The overall sequence variability in these two domains are very high compared to the other parts of the protein. The overall fold of 11 different modelled structures are similar however,at the ferredoxin inserted domain and at the C-terminal domain the same fold have been achievedby very different sequences. (**Figure 7**) This indicates two scenarios (a) the overall geometry achieved by the two different sequences somehow stabilising the fold or (b) both sequences have similar function but over a period of time their sequences have accumulated mutation due to variousselection pressure. Thus clearly indicating a case of divergent evolution of HdrA sequences.

**Figure 6:** Sequence alignment of HdrA sequences from 11 different clades are shown. The conserved residueare highlighted in yellow and the less conserved regions are shaded in green.

**Figure 7:**
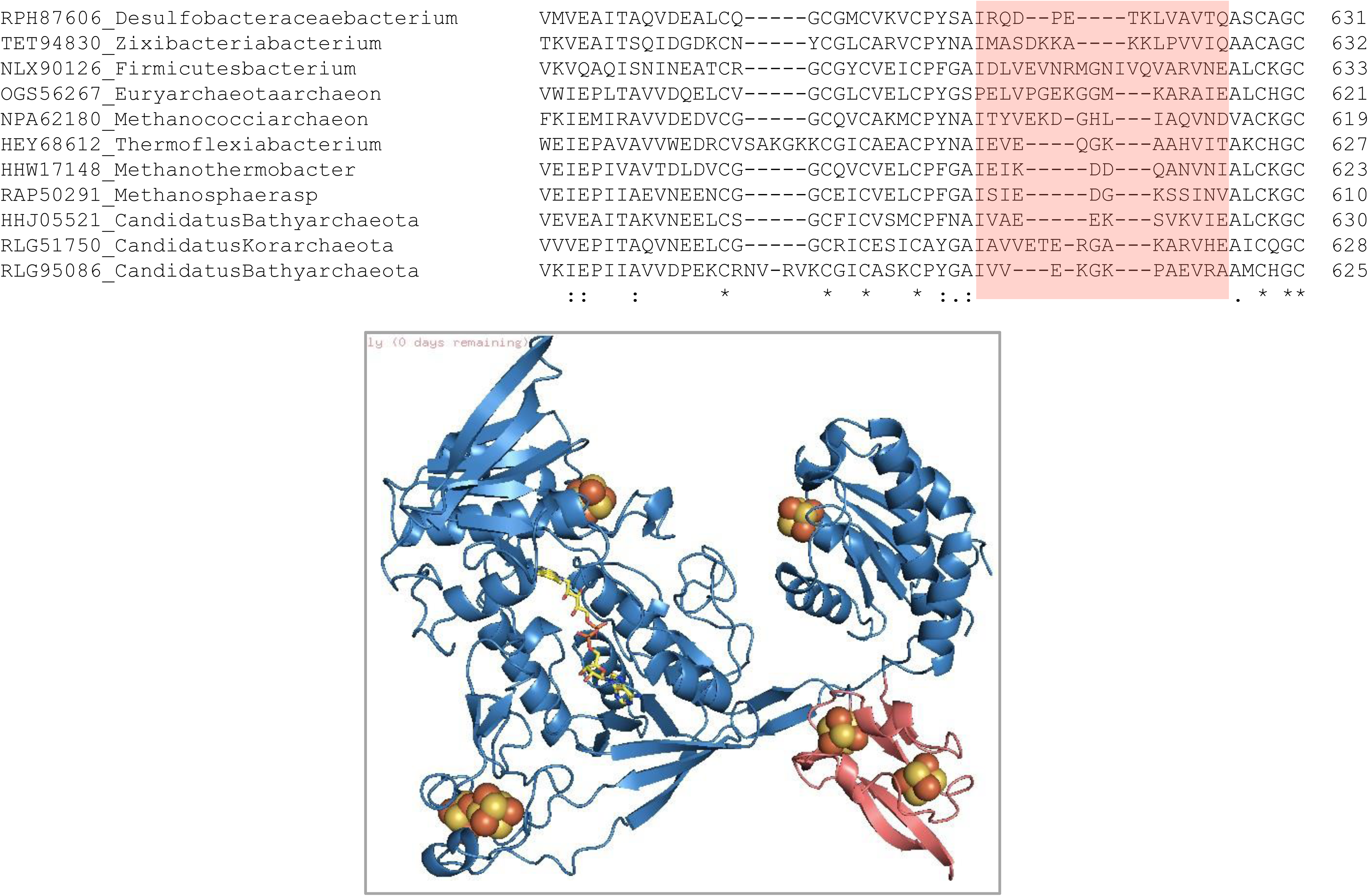
The variable domain in HdrA structure mapped. Inserted ferredoxin domain in green and C-terminalflexible domain in pink colour. The Fe-S causers are shown in red and yellow sphere.

### PPI network analysis; COGs and STRINGs

Table 2 summarises the PPI network data from STRING (version 11.5) (**37**) of 14 different homologous sequences, 11 of them are from 11 different clades and 3 more special sequences which are involved in ANME, nitrate fixation and sulphate metabolism.The interacting protein partners are very carefully chosen and the cutoff score was high (0.7) to exclude unwanted protein interactions. The 3rd column shows the names of individual proteins which are involved in the primary interaction with HdrA (**Table 2**). In most cases, the interactions of HdrA were observed with HdrB, HdrC, MvhD and MvhD subunits which are very obvious and crucial for the thiol-disulfidereduction reaction and electron bifurcation, the two critical steps of methanogenesis. However, for obligate methanogens HdrB, HdrC, MvhD and MvhD proteins are observed where as for non methanogenic organisms formate dehydrogenase, succinate dehydrogenase, oxidoreductases etc. proteins are observed. A complete shift in interacting partner is visible in Bathyarcheota BA1 and BA2; where the former showed interaction with acetyl CoA decarbonylase, PrcB_C domain containing protein, and sulphur carrier protein FdhD; whereas the later showed HdrA, HdrB, HdrC,sulfydrogenase and methyl enzyme M reductase. In Korarcheota Co-enzye F420 dehydrogenase and a very strong interaction was seen with CO dehydrogenase maturation factor. Firmicutes bacteria interacts with succinate dehydrogenase flavoprotein subunits, 2-polyprenylphenol hydroxyls and Uracil phosphoribosyltransferase interaction with a very high confidence score. For zixibacteria, interactions of Hdr with adenosylhomocysteine, succinate dehydrogenase, formate dehydrogenase, bifunctional homocysteine S-methyltransferase were visible. For desulfococcus multivorans HdrA interacts with other HdrB, HdrC, formate dehydrogenase, quinone interacting memerane bound oxide reducetase and pyridine nucleotide-disulfide reductase. Ammonifex degensii was found to interact with F420 non-reducing dehydrogenase and sulfydrogenase other than usual other HdrBand C subunits. In Methanoperedence the critical interactions are coneyme F420 hydrogenase, NADH ubiquinone oxidoreductase chain G-like protein with a very high confidence score. Similarly in Carboxxella the song interaction with the high confidence score are dehydrogenase, coneyme F420 hydrogenase and pyridine nucleotide diesel fide reductases other than heaterdisulfide reductase. In Syntropeceace bacterium the interacting partners are 4Fe-4S ferredoxin, succinate dehydrogenase and formate dehydrogenase among the critical ones. Ruminococcus champanellensis showed very unusual interactions with a reasonably high score and these are succinate dehydrogenatese/ fumarate reductase, uracil phosphoribosyl transferase and 2-propylphenol hydroxylase and related flavodoxin oxidoreductases.

## Discussion

### Homologous HDrA Sequences and phylogeny

HdrA the central subunit of HdrABC enzyme which carries out the critical reaction of thiol-disulfide reduction of the coenzymes (CoM and CoB, **38**) in methanogenesis and conserves energy in the same process carefully devised their functional role into three different subunits. Subunit A, (HDrA) the enzyme studied here, performs two critical functions; one is to bind flavin and other Fe-S clusters to bifurcate and transfer electron through those Fe-S clusters which is crucial for the redox reaction and the second one is it critically interacts with its binding partners to ferry the electrons to its proper destination (**39,40**). Collapse of any of these two functions makes the enzyme inactive and hence disrupting the pathway. The residues in HDrA sequence are carefully chosen it has 15 conserved residues out of 501 sequences and two CXXCXX motifs which binds to Fe-S cluster. Moreover, it has a very conserved flavin binding site with 17 conserved residues. The variable regions are observed between residues 235-315 and 568-654 where the sequences are <3% identical (**Figure 1**). It is observed throughout the study that the HDrA protein, which is known as methanogen specific is omnipresent. With a high sequence identity it is observed in Archaea and prokaryotic domain. The sequence identity with eukaryotes are very less and hence this report did not include them. These observations evokes two very fundamental questions; (i) if the organism is not engaging in methanogenesis then why it is preserving HdrA? (ii) whether HdrA is useful for the organism to perform some other function or it is using the protein in some other metabolic pathway? If yes then what are the evidences. The comparison of archaea and prokaryotic sequencesreveals that occasional insertion at the N and C terminal end for prokaryotes are visible (**Figure 1**) where for some archaea series deletion is observed. The overall sequence alignment suggests thatthe polyferrodoxin signature of HdrA, which is CXXCXX motif, is highly conserved in all sequences but most of the mutations were accumulated at the N and the C terminal end. The phylogenetic distances indicates the divergence of HDrA and confirms the HGT across species. Theparaphyletic distribution of HdrA across the phylogeny tree (**Figure 3**) among archaea and prokaryotes supports the horizontal transfer of HdrA gene. The multiple sequence alignment of all 501 sequences with minimum gap penalty and more conservation suggests a slow divergenceamong sequences which reflects in phylogeny as well. The alignment profile are more robust implying a less distant relationship among the sequences. The redox function is not limited to a single (methanogenesis) pathway rather they are distributed according to the need of organisms during evolution.

### Conserved domain Architecture and putative function

Investigation of coexisting domain were undertaken to understand the possible biological function of HdrA protein in archaeal and non-archaea homologues. 501 homologous sequences were distributed in 11 different domain architectures comprising both archaea and non-archaea suggests similar function of HdrA in these two kingdoms. Mapping the domain architecture on the phylogenetic tree (Figure 5) shows the conservative of core HDR domain in all species and the N and C terminal domain insertion and variation in different clades, suggesting the sequence diversification plays a greater role determining the domain architecture and finally the entirefunction of the protein. 11 different representative sequences from each group were chosen to generate individual models (SWISS MODEL) (**Figure 8**) showed very little overall structuralchanges among 11 different representative structures. The four key domains in the 3D structure of HDrA (the FAD binding domain, the ferredoxin insertion domain, the ferredoxin domain and the C-terminal domain) remains intact with little structural variation. The C-terminal variable region consists of pair of two beta sheets and two small helices and 4 loop regions (among which two are beta turns) which is significantly preserved in all the structures. (**Figure 7**) But the C-terminal sequences are extremely variable. The central beta sheet structure (at the C-terminal domain) with the turn is achieved by sequence stretches like; DPETK, DKKAKKL, EVNRMGNIVQVAR, PGEKGGMKAR, EKDGHLIAQ, EQGKAAH, IKDDQAN, SIEDGKSS, VAEEKSVK, VETERGAKAR, VVEKGKPAEV which does not explain directly the functional significance of this apparently variable region to preserve the fold so well by variety of sequences.

**Figure 8:**
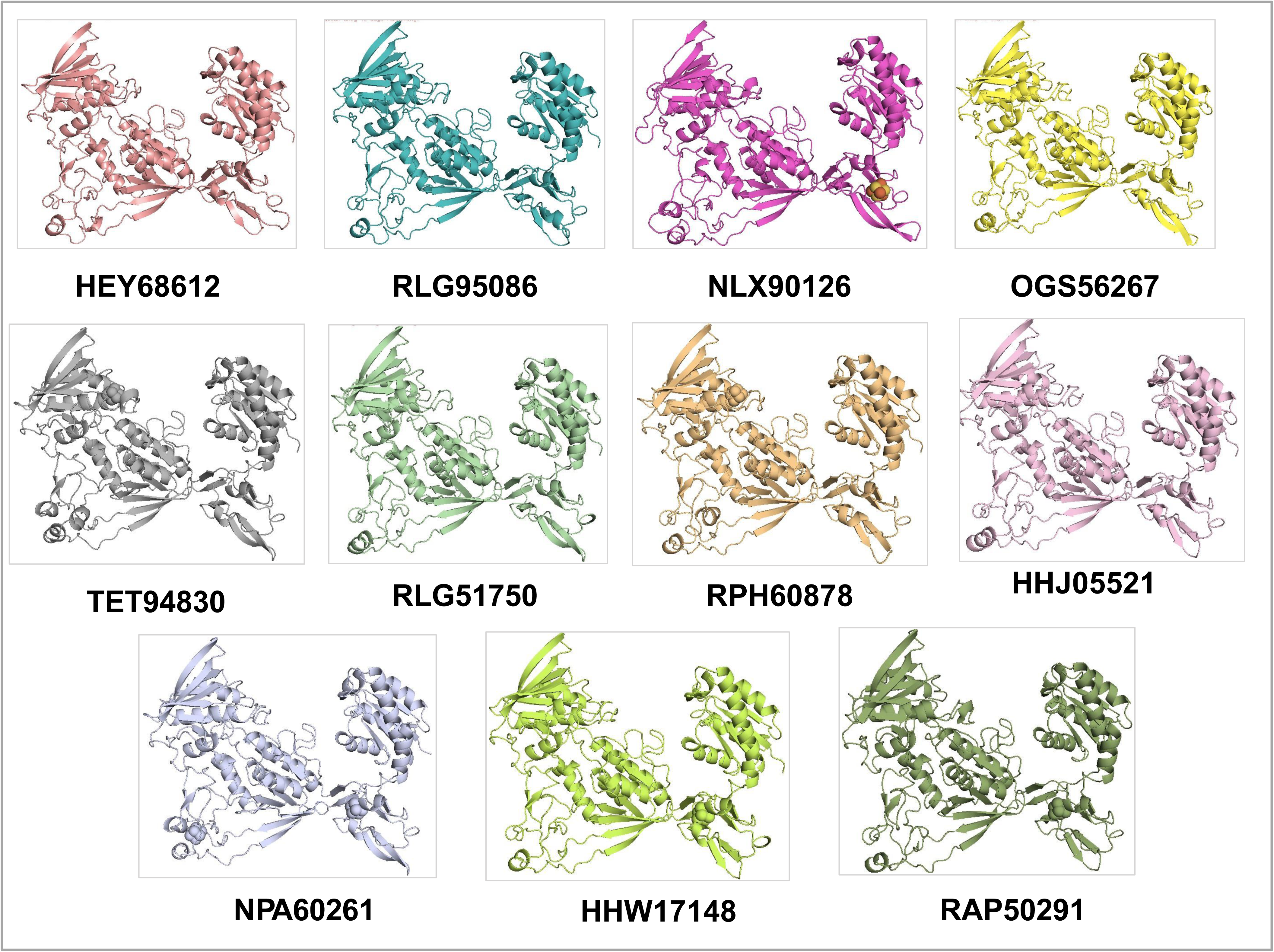
11 different HdrA structures are modelled in SWISS MODEL and shown.

### Homologous sequences and PPI network pattern

The key questions were asked here regarding HDrA function and evolution of HdrA function through phylogeny and structural analysis are, how HdrA adapt to the dynamic environment and how their evolution alters the protein interaction network? The sequence analysis of 11 different HdrA sequences and the PPI analysis using STRING database reveals how the sequences evolved and how the changes in sequences preserved the overall fold but changing the interacting partners in the network. To understand the interaction of PPI network which they achieved through changing physicochemical properties was throughly analysed using structural modelling of the homologous HdrA sequences. The variable domains; (i) Ferredoxin insertion domain and (ii) the C-terminal variable domains are least conserved (< 3%) regions in HdrA sequence. However, the sequence variability does not reflect on the structure as all of them holding the same fold. In divergent evolution the non-linear sequence-structure relationship is evident (**41**), where fitness contains prevent the formation of unstable intermediates during evolution (**42,43**). Thus preservation of secondary and tertiary structures despite sequence divergence are observed in various cellular proteins. In HdrA homologues, preservation of conserved domain as well as the variables domains inspite of sequence variability indicates either the timescale of sequence divergence is faster than the structural divergence or the the overall structure of the protein remained same to perform very similar functions but by altering the interacting partners or location which requires a substantial sequence alteration.

Figure 9 where PPI network represent a series of snapshots and the evolution of the enzyme and its interacting partners. The interaction network changes with species evolution which in turn directly correlates with their metabolic preferences. This also identifies a set of enzymes which are specific for specialised pathways in different organisms suggesting the common histories anddecent and lateral gene transfer. By correlating the structural features of the HDR homologues with PPI network and phylogenetic distribution highlights the the importance of environmental connection and metabolism among the organisms. Protein-protein interactions (PPIs) are one of the important components of biological networks and it is important to understand the evolutionary process of PPIs in order to understand how the evolution of biological networks has contributed to diversification of the existent organisms. Despite the significant role of molecular networks in determining an organism’s adaptation to its environment, we still do not know how such inter- and intra-molecular interactions within networks change over time and contribute to an organism’s evolvability while maintaining overall network functions. Even though the main structure and function of an essential network may be conserved among divergent species over history, the individual components of the network may exhibit divergent properties. Thus functionally divergentproperties may be observed at the molecular level despite selection maintaining structurally conserved properties at the network level (**44–48**). Figure 10 mapped the individual network on the species specific phylogenetic tree suggesting the evolution of the individual network and directly correlating the metabolic diversification during evolution.

**Figure 9:**
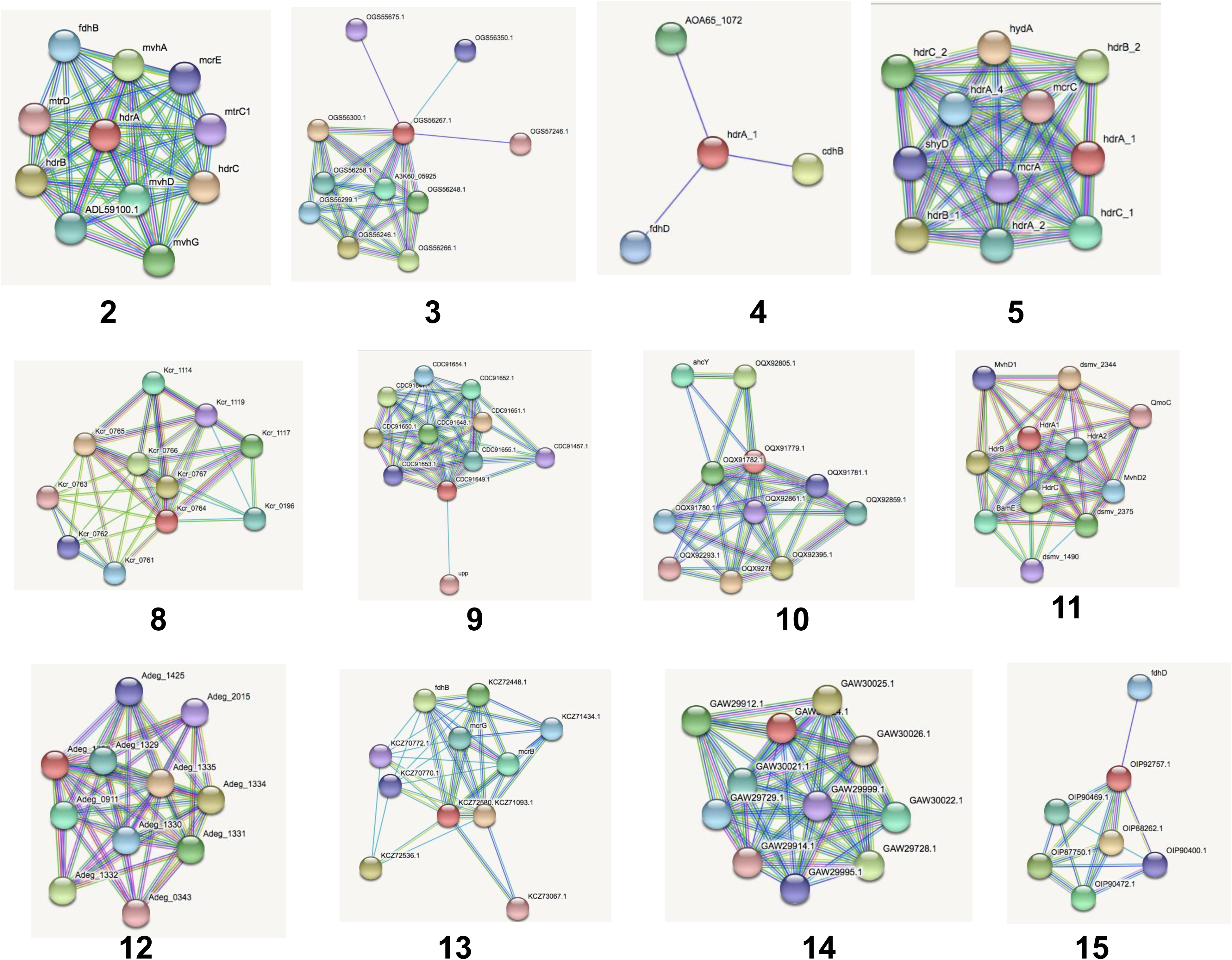
Protein protein interaction network of HdrA homologues are shown. The network was modelledusing STRING 11.5. The interaction score was set >0.7 to construct individual PPI network.

**Figure 10:**
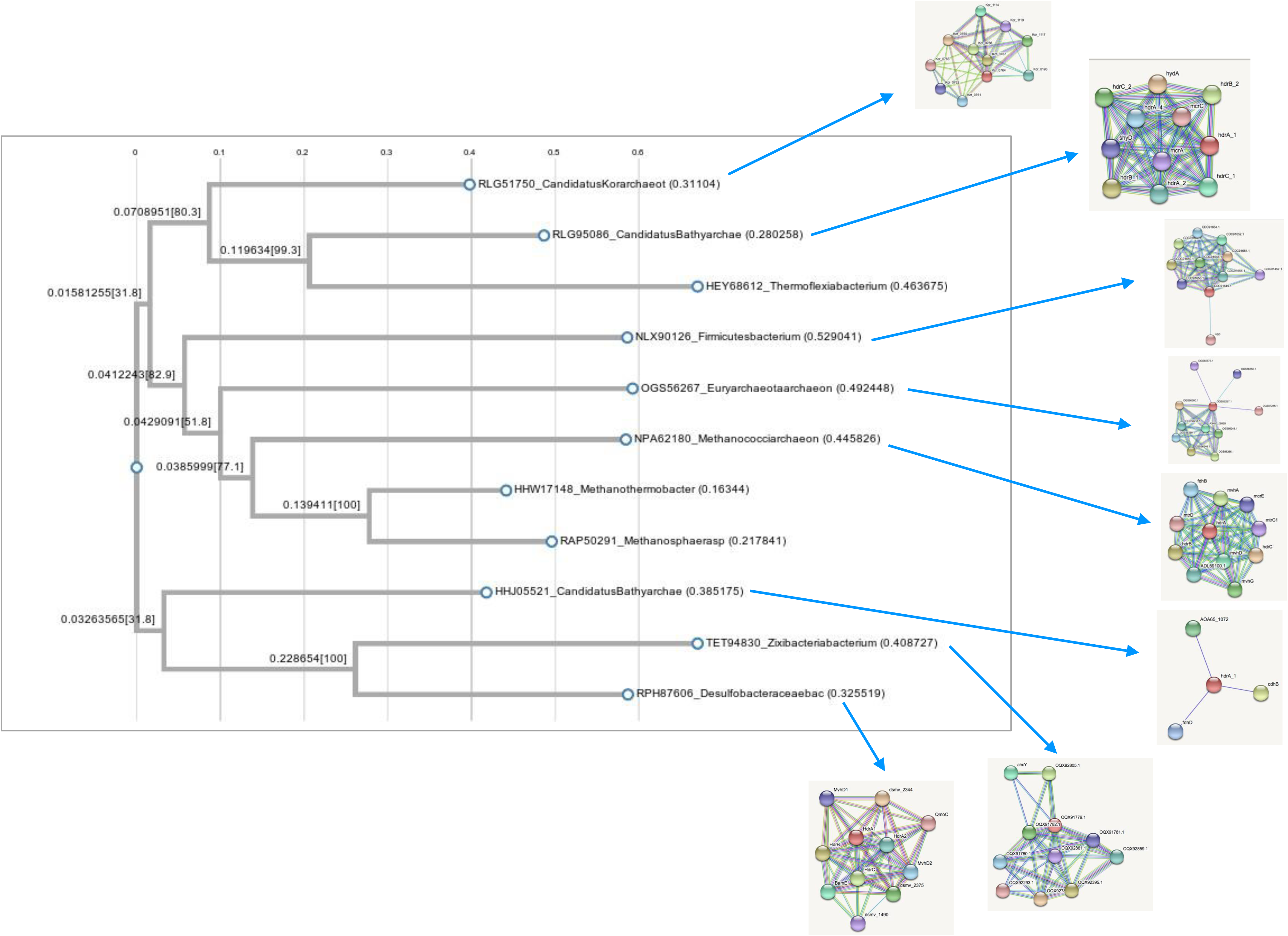
PPI network mapped on phylogeny tree.

## Conclusion

The sequence analysis and phylogeny provides the first insight into an evolutionarily adapted redox protein which can also participate in electron transfer, happens to work with both anaerobic and aerobic organism. It suggests that the basic mechanistic principle of redox reaction with multiple Fe-S cluster disulphide, sulphate reduction, nitrate oxidation use similar mechanistic approach. The sequence analysis, phylogeny and domain architecture analysis of HDrA sequences from bacterial and archaea sources reflects the broad difference in their metabolism and its evolution during various changes in the environment. Environmental adaption and metabolic changes were inherited through accumulation of meaningful mutation at the protein level which eventually facilitate the functional divergence and speciation. This study provides a robust analysis of linear sequences and domain architecture which suggests the evolution of molecular sequences along with structural domain changes the functional adaptation for different metabolic processes by just repurposing the existing enzyme with minimum sequence variation.

## Abbreviation

Mj: Methanococcus jannaschii
Hdr: Hetero disulphide reductase
FBEB: Flavin Based Electron Bifurcation
FMN: Flavin mono-nucleotide
FAD: Flavin di-nucleotide

## Acknowledgements

This work is supported by DST-WOS-A Scheme (project no-SR/WOS-A/LS-70/2018) and partially by DBT Biocare (project no BT/PR31884/BIC/101/1212/2019) The author would like to thank Prof. K. N Ganesh as an institute mentor and Indian Institute of Science Education and Research - Tirupati (IISER - Tirupati, India) for hosting and infrastructural support.

## Conflict of interest

There is no conflict of interest.

## Notes

### Competing Interest Statement

The authors have declared no competing interest.

